# Tree-aware conditional language modeling recovers mutational patterns of viral evolution

**DOI:** 10.64898/2026.08.25.746971

**Authors:** Polina V Polunina, Wolfgang Maier, Alan F Rubin

## Abstract

The evolutionary accessibility of a protein mutation depends on the sequence background in which it arises and its lineage history, yet most protein language models estimate sequence plausibility without explicitly considering the ordered sequence changes through which descendants arise. We developed evoPLM-Tree, a tree-aware conditional autoregressive language model that predicts descendant protein sequences from ancestral sequences together with phylogenetically derived evolutionary features. We demonstrated our approach using SARS-CoV-2 spike protein, pairing sequences from early Omicron lineages according to their positions on a mutation-annotated phylogeny, and evaluating model performance on sequence pairs from later lineages. Prompt-masking experiments showed that incorporating phylogenetic context substantially increased reliance on the supplied input information compared with a sequence-only model. Generated descendant sequences accurately reproduced the positional distribution of mutations observed during viral evolution, with strong correlations between predicted and observed mutation-frequency profiles for both the receptor-binding domain (Spearman’s *ρ* = 0.823) and the full spike protein (*ρ* = 0.736). Although prediction accuracy for individual substitutions decreased with increasing evolutionary distance, the model consistently captured aggregate mutational patterns across the spike protein. Model-assigned mutation probabilities were also enriched among substitutions experimentally tolerated in deep mutational scanning assays of Omicron BA.2 receptor-binding domain expression (1.19-fold enrichment) and ACE2 binding (1.04-fold enrichment), despite the model being trained solely on observed ancestor–descendant sequence pairs and associated phylogenetic context features. These results demonstrate that explicitly providing protein language models with phylogenetic context during sequence generation can recover lineage-specific mutational patterns and yields probabilistic predictions consistent with experimentally measured functional constraints. evoPLM-Tree provides a framework for modeling protein evolution along phylogenetic lineages and prioritizing plausible future mutations from genomic surveillance data.

**Author summary:** Viruses evolve by accumulating mutations along branching lineages, so the changes a virus is likely to acquire next depend on its current sequence. Computational models that predict protein sequences usually ignore this history: they judge whether a sequence looks plausible in general, not whether it is a plausible descendant of a particular ancestor. We asked whether giving a language model explicit information about evolutionary history would help it reproduce how a viral protein actually changes. Using the SARS-CoV-2 spike protein, for which millions of real sequences have been arranged into a detailed evolutionary tree, we trained a model on pairs of ancestor and descendant sequences together with simple measurements taken from the tree, such as how many mutational steps separate the two. Our resulting model, evoPLM-Tree, benefitted substantially more on the information it was given than a model trained on sequences alone, reproduced where mutations occur across spike, and assigned higher probability to substitutions that laboratory experiments show the protein tolerates. As surveillance data are captured for other pathogens, the same approach could help anticipate mutations in newly emerging viruses.

## Introduction

Protein evolution is a sequential process in which mutations accumulate along branching lineages. Which substitutions are observed at a given point in a lineage therefore depends not only on their biochemical and fitness effects but also on the substitutions already present in the same protein and on the evolutionary path by which that sequence was reached. The effects of most substitutions in a protein combine approximately additively, and strong interactions between sites are the exception rather than the rule. Path dependence nonetheless arises for a simpler reason: a substitution can only be observed on the sequence background that a lineage has actually reached, so the set of accessible next steps changes as a lineage accumulates changes. Ongoing surveillance of SARS-CoV-2 provides a unique opportunity for investigating these processes because millions of viral genomes have been collected and organized into temporally and phylogenetically resolved lineages. Its spike glycoprotein is a major target of such analyses because it mediates entry through the host ACE2 receptor and is exposed to strong selective pressures associated with receptor usage and antibody-mediated immunity [1].

The receptor-binding domain (RBD) of spike illustrates why viral mutation effects cannot be evaluated independently. Experimental data generated using deep mutational scanning (DMS) has shown that amino-acid substitutions differ widely in their effects on RBD expression, ACE2 binding, and antibody recognition, and that these effects can shift between SARS-CoV-2 variant backgrounds, such as the ancestral and Omicron RBDs [2, 3]. More recently, full-spike DMS demonstrated that experimentally measured effects on ACE2 binding, cell entry, and antibody escape can help explain the evolutionary success of SARS-CoV-2 lineages [4]. In the Omicron BA.1 lineage, combinations of affinity-enhancing substitutions compensate for the deleterious effects of several antibody-escape mutations, allowing substantial antigenic change while maintaining ACE2 binding [5]. However, the effects of most substitutions are broadly conserved across variant RBD backgrounds, with large shifts restricted to a minority of mutations [3]. These observations show that the evolutionary accessibility of a mutation depends partly on the substitutions already present in the lineage. Models capable of identifying such accessible changes can help prioritize variants for experimental characterization and connect population-level genomic surveillance with molecular measurements of protein function [4].

Epidemiological models use changes in lineage prevalence across time and geographic regions to estimate viral fitness. For example, PyR0 applied a hierarchical Bayesian model to millions of SARS-CoV-2 genomes to estimate lineage growth rates and mutation-associated fitness effects, enabling emerging lineages to be ranked from their mutational profiles [6]. Such approaches make effective use of surveillance data and quantify epidemiological success, but their primary outputs are lineage growth or substitution-associated fitness rather than probabilities of a complete protein sequence arising from a given ancestor.

Statistical evolutionary models instead characterize constraints and mutability from sequence variation, substitution processes, or interactions among sites. Models incorporating temporal occurrence, structural annotations, and mutation frequencies have been used to estimate which SARS-CoV-2 substitutions may emerge in subsequent variants. Epistatic sequence models trained on coronavirus homologs have also predicted mutable and constrained positions more accurately than independent-site conservation models, demonstrating that correlations between residues contain information about evolutionary accessibility [7]. These methods generally assign scores to mutations, sites, or complete sequences relative to a reference, but do not directly learn to generate a descendant sequence from its ancestral sequence.

Generative language models learn distributions over protein sequences and can therefore represent combinations of mutations rather than scoring only predefined substitutions. Hie et al. [8] used language-model grammaticality and semantic change to identify mutations that remained compatible with viral sequence constraints while potentially enabling immune escape in influenza, HIV, and SARS-CoV-2. GenSLMs extended autoregressive modeling to complete viral genomes, using large-scale pretraining and SARS-CoV-2 fine-tuning to learn representations that distinguished variants and reflected pandemic genomic diversity [9]. Kumar [10] used an encoder–decoder model to translate spike sequences between adjacent SARS-CoV-2 clades [11] and reproduce clade-level substitution patterns. PandoGen was developed to generate complete future SARS-CoV-2 spike sequences from surveillance data, tailoring model training toward prospective sequence forecasting [12]. These models establish the utility of generative sequence modeling, but they estimate sequence plausibility, clade-level transformations, or future population-level sequence distributions without training on individual ancestor–descendant sequence pairs extracted from a densely resolved phylogeny.

Temporal forecasting methods explicitly exploit changes in sequence distributions over successive periods. Ma et al. [13] constructed population-level mutation frameworks from temporally grouped Omicron S1 sequences, sampled amino-acid combinations using empirical mutation profiles, and evaluated candidate variants against later calendar intervals. Their approach successfully incorporated both recurrent mutational structure and stochastic variation, but it operated without a phylogenetic tree and did not anchor predictions to a particular ancestral sequence. Thus, temporal separation alone does not explicitly represent the lineage-specific paths through which descendants arise.

Pretrained protein language models provide complementary estimates of biochemical and evolutionary constraint learned from broad protein sequence corpora. Their likelihoods support zero-shot mutation-effect prediction [14], while their embeddings represent structural and functional relationships. Evo-velocity used local likelihood differences from a protein language model to infer directional vector fields in sequence-representation space and recover evolutionary trajectories across diverse protein families [15]. More recently, Lamb et al. [16] applied the pretrained ESM-2 model [17] to SARS-CoV-2 spike, showing that its likelihoods recover regional constraints, support in-silico mutational analysis, and encode relationships among observed variants and trajectories. Other protein language models incorporate evolutionary information more directly through homologous sequences: MSA Transformer [18] jointly models aligned sets of homologs, while autoregressive approaches such as PoET [19] and ProtMamba [20] condition sequence modeling or generation on collections of related proteins, and Tranception [21] can incorporate homologous sequences retrieved at inference. More recently, Phyla introduced an explicit tree-based training objective for learning phylogenetic relationships among protein sequences [22]. These approaches show that evolutionary information beyond an individual sequence can improve protein representation, generation, and mutation-effect prediction.

A remaining gap is the direct modeling of viral protein evolution from observed ancestor–descendant sequences on a phylogenetic tree. In this formulation, the target is not the plausibility (or inferred fitness) of an individual sequence, but the conditional probability of a descendant sequence given a defined ancestor. Existing approaches have not directly trained autoregressive protein sequence models on individual ancestor–descendant sequence pairs while jointly conditioning descendant generation on such tree-derived features.

We developed evoPLM-Tree (Fig 1A-D, S1 Fig) to distinguish general protein-level constraints from lineage- and background-dependent effects while retaining the capacity to generate complete descendant sequences. evoPLM-Tree is a tree-aware conditional language model that learns to predict descendant protein sequences from ancestor–descendant sequence pairs extracted from a mutation-annotated phylogeny, conditioned on a related starting sequence and tree-derived evolutionary context. We demonstrate this approach on SARS-CoV-2 spike protein, which provides an unusually dense and phylogenetically resolved dataset for evaluating whether such a model can recover biologically meaningful mutational patterns. We show that our formulation reproduces the positional distribution of mutations across the spike protein and assigns greater probability to substitutions that retain function in DMS experiments. Through these analyses, we determine whether explicitly learning from viral evolutionary history improves prediction of protein function.

**Fig 1.**
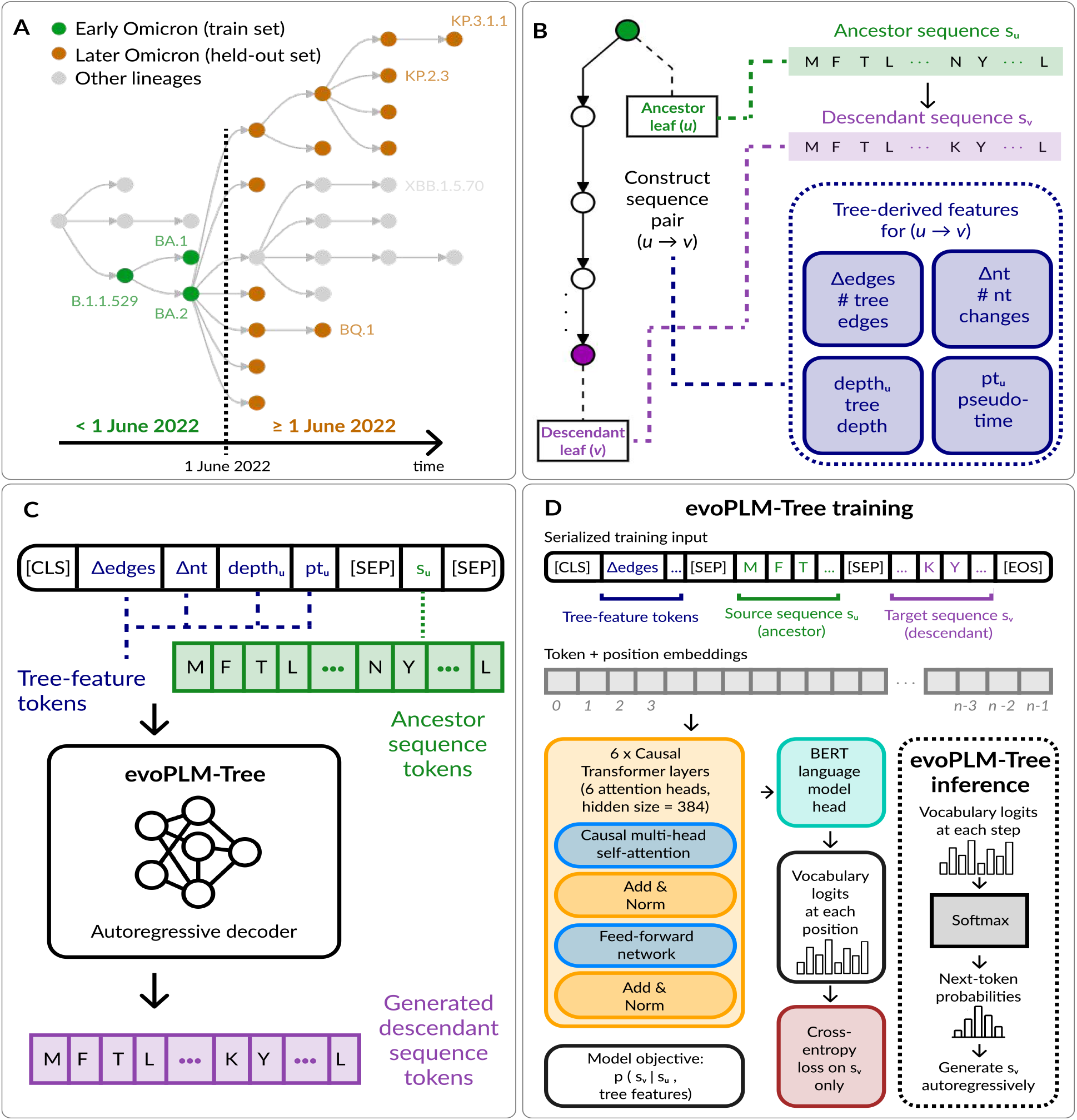
Overview of evoPLM-Tree. **(A)** SARS-CoV-2 spike sequences were mapped to the UShER mutation-annotated tree (MAT) and split at 1 June 2022 into training and held-out sets. **(B)** For each phylogenetic sequence pair (*u* → *v*), the ancestral sequence *s*_*u*_, descendant sequence *s*_*v*_, and tree-derived features describing the relationship between *u* and *v* comprised the number of intervening tree edges (Δ_edges_), nucleotide changes (Δ_nt_), ancestor tree depth (depth_*u*_), and ancestor mutational pseudotime (pt_*u*_) were extracted. **(C)** evoPLM-Tree conditions descendant-sequence generation on the ancestral sequence and tree-derived phylogenetic features. These inputs provide the evolutionary context used by the autoregressive decoder to generate descendant-sequence tokens. **(D)** During training, tree-feature tokens, *s*_*u*_, and *s*_*v*_ are processed by a six-layer causal Transformer followed by a BERT language-model head. Cross-entropy loss is computed only over *s*_*v*_. During inference, the model autoregressively generates the descendant sequence, learning *p*(*s*_*v*_ | *s*_*u*_, tree features).

## Results

### evoPLM-Tree performance depends on evolutionary conditioning context

We trained evoPLM-Tree on pairs of ancestor and descendant sequences from a mutation-annotated SARS-CoV-2 phylogeny [23]. Separate models were trained for the spike receptor binding domain (RBD) and the full spike protein using data from early Omicron lineages collected before 1 June 2022. Performance was evaluated on sequence pairs from later non-recombinant Omicron lineages. After removal of trivial source–target pairs (i.e., pairs with identical source and target protein sequences or no increase in root-to-tip nucleotide mutational pseudotime), the training datasets contained 87,370 RBD sequence pairs and 180,079 full-spike sequence pairs, whereas the held-out datasets contained 272,133 RBD sequence pairs and 520,091 full-spike sequence pairs (see Materials and methods).

To determine whether the models relied on their conditioning context, we compared two variants with the same transformer architecture: a sequence-only model, conditioned on (i.e. prompted with) only the starting sequence, and a tree-aware model (evoPLM-Tree), conditioned on the starting sequence together with tree-derived evolutionary features. We evaluated the observed descendant sequences under teacher forcing while either retaining or masking the complete conditioning prompt (see Materials and methods). Masking the prompt decreased performance for all models (Fig 2A,B). For the RBD, prompt masking increased the loss by 4.8% (loss ratio 1.048) in the sequence-only model and by 24.2% (loss ratio 1.242) in the tree-aware model (Fig 2A). For full spike, the corresponding loss increases were 27.5% (loss ratio 1.275) and 73.6% (loss ratio 1.736) (Fig 2B). Thus, all models used information from the conditioning prompt, but prompt dependence was consistently greater for the tree-aware models and was strongest for full-spike prediction.

**Fig 2.**
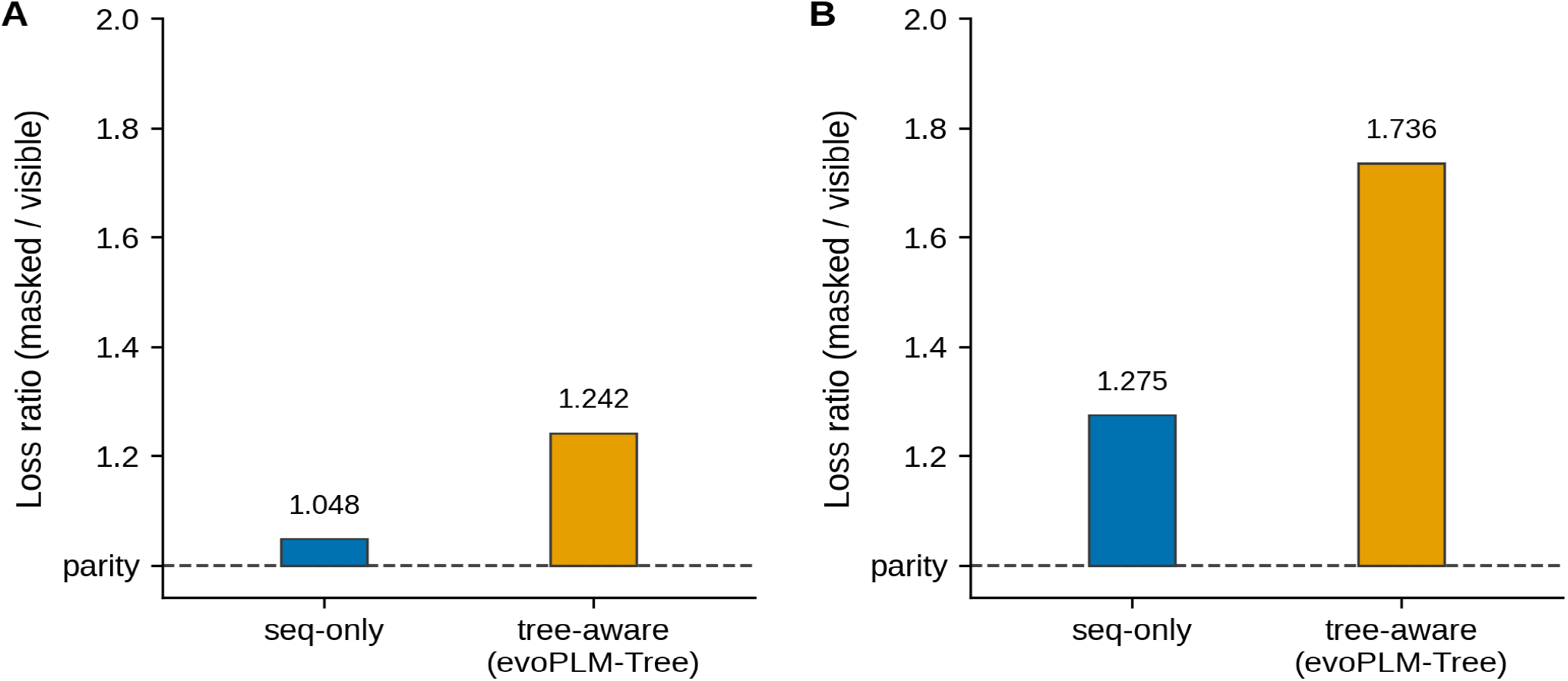
Prompt dependence of sequence-only and tree-aware models for (A) RBD and (B) full spike protein. For both panels, prompt dependence was quantified as the ratio of descendant-token teacher-forced cross-entropy with the conditioning prompt masked to that with the prompt visible for the sequence-only and tree-aware (evoPLM-Tree) models. The dashed line denotes parity, indicating no effect of prompt masking; values above parity indicate increased prediction loss when the conditioning context is removed. **(A)** For the RBD, masking increased the loss ratio to 1.048 for the sequence-only model and 1.242 for the tree-aware model. **(B)** For full spike, the corresponding ratios were 1.275 and 1.736. In both tasks, the larger increase for evoPLM-Tree indicates greater dependence on the conditioning context.

The larger effect of prompt masking in the tree-aware model was observed for both prediction tasks, with the difference between tree-aware and sequence-only loss ratios increasing from 0.194 for the RBD to 0.461 for full spike. These results indicate that the tree-aware model makes greater use of its conditioning context when predicting descendant sequences.

### Generated RBD sequences reproduce observed mutation hotspots

We next asked whether evoPLM-Tree captured the positional distribution of mutations observed during SARS-CoV-2 evolution. Using the RBD-trained evoPLM-Tree model, positional mutation-frequency profiles were computed for observed and model-generated descendants from 2,000 held-out sequence pairs (see Materials and methods).

The predicted positional mutation-frequency profile closely followed that observed in the held-out sequence pairs (Fig 3A, S1 Table). Across the 223 RBD positions, observed and predicted mutation frequencies were strongly correlated (Spearman’s *ρ* = 0.823, two-sided *p* = 2.90 *×* 10^−56^; Fig 3C, S1 Table).

**Fig 3.**
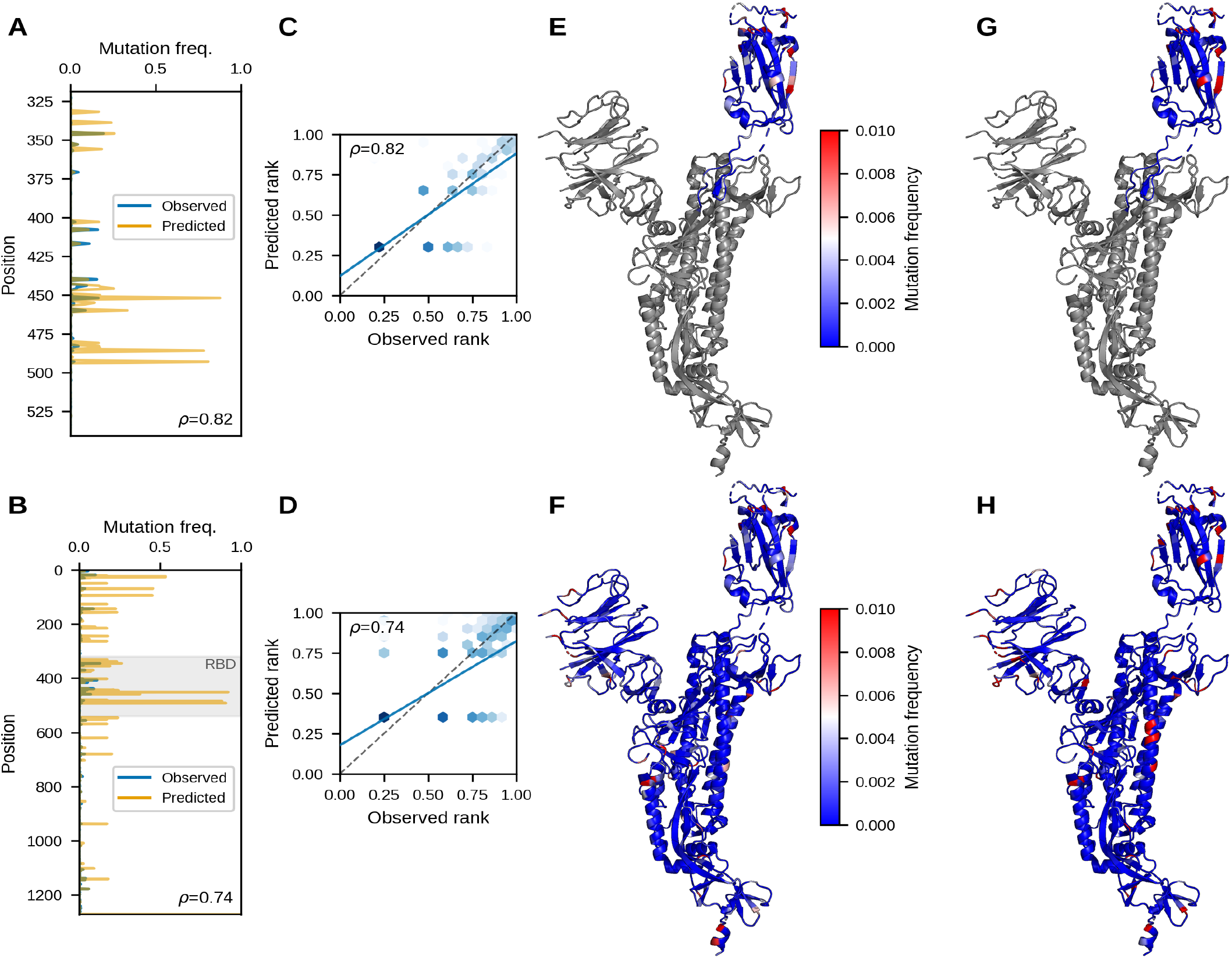
evoPLM-Tree captures the positional and structural landscape of SARS-CoV-2 spike mutations. **(A-B)** Observed (blue) and predicted (orange) positional mutation frequencies from 2,000 stratified held-out pairs, each panel using the correspondingly trained model. **(A)** RBD residues 319–541 (223 positions), Spearman *ρ* = 0.823 (two-sided *p* = 2.90 *×* 10^−56^). **(B)** Full spike (1,274 positions, zero-based 0–1273), *ρ* = 0.736 (*p* = 4.11 *×* 10^−218^); the gray band marks the RBD (residues 319–541). In-panel *ρ* values are rounded to two decimals. **(C-D)** Rank agreement for the same data, with frequencies converted to percentile ranks; darker hexagons denote more positions per bin, scaled independently per panel. The solid blue line shows the linear regression fit and the dashed diagonal denotes perfect agreement. **(C)** RBD. **(D)** Full spike. **(E-H)** Per-residue frequencies projected onto chain A of the spike structure (PDB 6VSB), colored blue (0) through white (0.005) to red (≥ 0.01, saturated): observed **(E, F)** and evoPLM-Tree-predicted **(G, H)** for the RBD **(E, G)** and full spike **(F, H)**. For RBD panels, regions outside residues 319–541 are gray; for full-spike panels, only residues resolved in the experimental structure are shown.

Positions that mutated frequently in the observed descendants also tended to receive high mutation frequencies in model-generated sequences, whereas more constrained positions remained comparatively conserved. The agreement was not limited to a small number of individual residues but extended across the overall RBD mutation-frequency profile.

Projection of the positional frequencies onto the spike structure showed that observed and predicted mutations were concentrated in similar surface-exposed regions of the RBD (Fig 3E,G, S3 Table). Elevated frequencies were particularly apparent within the receptor-binding motif, which contains residues directly involved in ACE2 interaction and major targets of antibody-mediated selection [24]. evoPLM-Tree therefore recovered both individual high-frequency sites and the broader spatial organization of the RBD mutational landscape.

The generated sequences showed higher overall mutation frequencies than the observed descendants (Fig 3A, S1 Table). This difference affected the magnitude of predicted mutability more strongly than its positional distribution, because the principal mutational hotspots remained concordant between observed and generated sequences.

### evoPLM-Tree recovers the mutational landscape of full spike

We repeated the positional mutation-frequency analysis using the full-spike-trained evoPLM-Tree model, generating descendants from 2,000 stratified held-out sequence pairs (see Materials and methods). The full spike sequence comprised approximately 1,274 amino acids in the evaluated alignment, compared with 223 amino acids for the RBD. Modeling full spike is therefore more challenging because the sequence is substantially longer and contains multiple domains subject to distinct structural, functional, and immunological constraints.

Nevertheless, the model-generated mutation-frequency profile remained strongly associated with the profile observed in held-out sequence pairs (Fig 3B, S2 Table). Across the full spike sequence, observed and predicted positional mutation frequencies were strongly correlated (Spearman’s *ρ* = 0.736, two-sided *p* = 4.11 *×* 10^−218^; Fig 3D, S2 Table). Predicted mutations were concentrated in many of the same regions as naturally observed substitutions (Fig 3F,H, S4 Table). These included the receptor-binding domain, the N-terminal domain, and exposed loop regions in the spike ectodomain. In contrast, more structurally constrained regions generally exhibited lower predicted and observed mutation frequencies.

The lower correlation for full spike than for the RBD likely reflects the greater sequence length and the diversity of constraints operating across distinct spike domains. However, the persistence of a strong genome-wide positional association indicates that evoPLM-Tree learned a structured mutational landscape rather than reproducing a small set of recurrent RBD substitutions.

### Exact mutation recovery varies with evolutionary distance

Recovering the positional distribution of mutations does not necessarily mean that a model predicts the exact amino-acid substitutions present in an individual descendant sequence. We therefore measured exact mutation recovery as the fraction of observed amino-acid substitutions between the input and descendant sequences that were reproduced exactly in the model prediction. This analysis used 2,000 observed input–descendant sequence pairs together with evoPLM-Tree-generated descendants for the corresponding input sequences. A substitution was considered recovered only when its position, ancestral amino acid, and descendant amino acid all matched. For each mutation-count bin, the reported value is the mean of this per-example recovery fraction (see Materials and methods).

For the RBD model, mean exact mutation recovery was highest for sequence pairs containing one mutation, at 0.45 (the number of evaluated sequence pairs contributing to this mutation-count bin *n* = 1,192). Recovery was 0.32 for two mutations (*n* = 527) and for three mutations (*n* = 212), 0.26 for four to five mutations (*n* = 66), and 0.15 for six to ten mutations (*n* = 3; Fig 4A, S1 Table). No RBD examples contained 11 or more mutations. Thus, RBD recovery was generally lower at greater evolutionary distances, although the change was not strictly monotonic and the estimate for six to ten mutations was based on very few examples.

**Fig 4.**
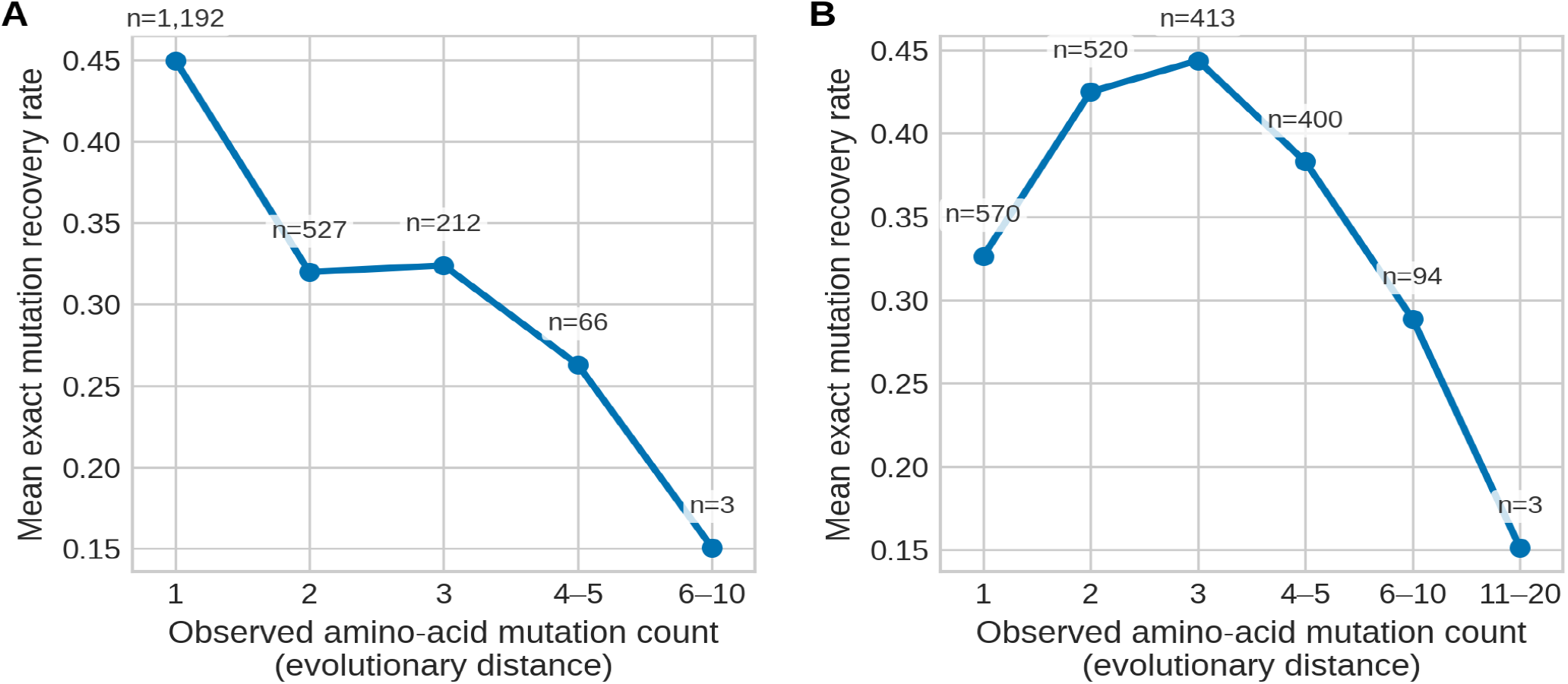
Exact mutation recovery as a function of evolutionary distance for (A) RBD and (B) full-spike. Mean exact mutation recovery was calculated for 2,000 observed input–descendant sequence pairs grouped by the number of amino-acid mutations separating each pair. For each pair, recovery was defined as the fraction of observed mutations reproduced exactly by evoPLM-Tree, requiring agreement in mutation position, ancestral residue, and descendant residue. Points show bin means, connecting lines are guides to the eye, and n denotes the number of sequence pairs in each bin.

A different pattern was observed for the full-spike model. Mean recovery was 0.33 for one mutation (*n* = 570), increased to 0.43 for two mutations (*n* = 520), and reached its maximum of 0.44 for three mutations (*n* = 413). It subsequently decreased to 0.38 for four to five mutations (*n* = 400), 0.29 for six to ten mutations (*n* = 94), and 0.15 for 11–20 mutations (*n* = 3; Fig 4B, S2 Table). Exact recovery therefore peaked at an intermediate evolutionary distance for the spike model rather than at the shortest distance. Again, the estimate for the largest-distance bin should be interpreted cautiously because it was based on only three examples.

The mutations themselves likely differ with distance: across full spike, single-change pairs are dominated by sporadic substitutions, whereas two- and three-change pairs more often correspond to recurrent, lineage-defining combinations seen repeatedly during training. RBD substitutions are already concentrated at a few recurrent antigenic positions, so even single-mutation pairs are enriched for predictable changes, consistent with the higher recovery at one mutation for RBD (0.45) than for full spike (0.33).

Overall, exact substitution recovery was lower for larger mutational steps after the model-specific peak: one mutation for RBD and three mutations for spike. This pattern indicates that the number of possible descendant sequences grows with evolutionary distance. However, the sparsely populated largest-distance bins prevent strong conclusions about performance at the greatest evolutionary distances. Together with the preceding positional analyses, these results suggest that evoPLM-Tree can capture aggregate mutational tendencies even when it does not reproduce every substitution in a particular descendant sequence.

### Model probabilities are enriched for experimentally tolerated RBD mutations

To determine whether evoPLM-Tree probabilities reflected experimentally measured functional constraints, we compared mutation probabilities assigned by the RBD-trained evoPLM-Tree model with deep mutational scanning measurements for the Omicron BA.2 RBD reported by Starr et al. [25] (see Materials and methods). evoPLM-Tree preferentially assigned probability to substitutions that retained experimentally measured RBD function.

For RBD expression, the probability-weighted enrichment was 1.19, indicating that the model assigned 19% more probability mass to tolerated mutations than expected from their background frequency in the evaluated mutation set. For ACE2 binding, the corresponding enrichment was 1.04, representing a 4% increase over random expectation (Fig 5, S5 Table– S7 Table).

**Fig 5.**
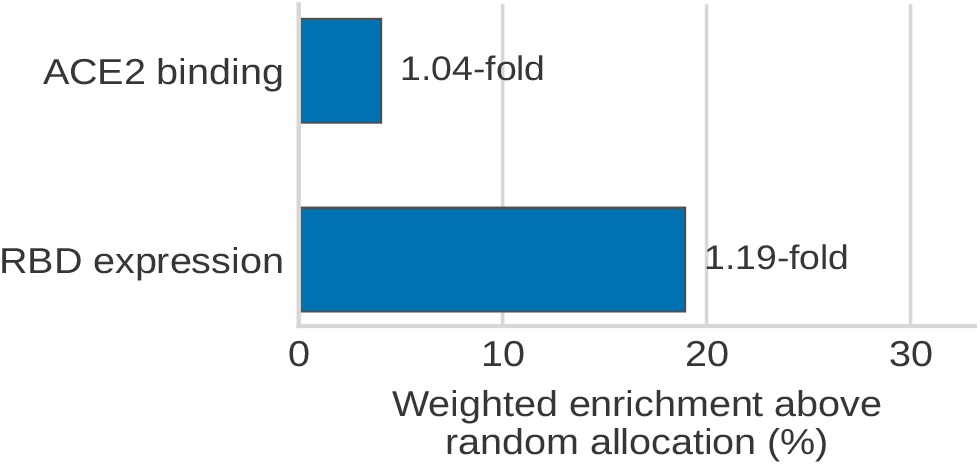
Enrichment of experimentally tolerated BA.2 RBD mutations among evoPLM-Tree predictions. Model probabilities for 3,801 single amino-acid substitutions were compared with deep mutational scanning measurements of RBD expression (`urn:mavedb:00000671-a-1`) and ACE2 binding (`urn:mavedb:00000670-a-1`) from MaveDB. Mutations with DMS scores greater than or equal to zero were classified as tolerated. Weighted enrichment was calculated as the model-probability-weighted fraction of tolerated substitutions divided by their unweighted fraction among all evaluated substitutions. Bars show this quantity expressed as the percentage increase over random allocation of probability mass, that is (*E*_weighted_ − 1) *×* 100; zero therefore corresponds to the uniform-allocation baseline and positive values indicate preferential allocation of model probability to experimentally tolerated mutations. Enrichment was 1.19-fold (19% above baseline) for RBD expression and 1.04-fold (4% above baseline) for ACE2 binding.

The stronger enrichment for RBD expression suggests that model probabilities more clearly reflected constraints associated with protein expression or folding than constraints associated specifically with ACE2 interaction. Importantly, the model was trained only on observed ancestor–descendant sequence pairs and associated phylogenetic context features and was not optimized using the DMS measurements. The enrichment therefore indicates that the learned evolutionary distribution contains information associated with experimentally measured functional tolerance.

## Discussion

We developed evoPLM-Tree, a tree-aware conditional language model that learns to predict descendant protein sequences from ancestor–descendant sequence pairs, conditioned jointly on an ancestral sequence and phylogenetically derived evolutionary context. Rather than modeling sequence plausibility or population-level sequence distributions alone, as in several previous applications of language models to viral evolution [8–10, 12], evoPLM-Tree explicitly learns from ancestor–descendant sequence pairs extracted from a mutation-annotated phylogeny. We demonstrated this approach on SARS-CoV-2 spike, which offers the dense genomic surveillance and phylogenetically resolved sequences needed to train and rigorously evaluate such a model. Across held-out Omicron lineages, the model reproduced the positional distribution of naturally occurring mutations, assigned increased probability to experimentally tolerated substitutions, and showed substantially greater reliance on evolutionary conditioning than a sequence-only model. Together, these results demonstrate that incorporating phylogenetic information into conditional sequence generation enables language models to recover biologically meaningful patterns of viral protein evolution.

A central observation of this study is that phylogenetic context contributed measurable predictive information beyond the ancestral sequence itself. Prompt-masking experiments showed that removing the conditioning context increased prediction loss for both sequence-only and tree-aware models, but the effect was consistently larger for evoPLM-Tree, particularly for full-spike prediction. These findings indicate that the additional tree-derived features were actively used during inference rather than ignored by the model. Because the training objective required prediction of observed descendants from specific ancestral backgrounds, the model learned conditional evolutionary distributions that depend not only on protein sequence constraints but also on lineage-specific evolutionary history. This is consistent with experimental evidence that the accessibility of SARS-CoV-2 substitutions depends on the spike sequence background on which they arise [3, 5], although the model conditions on the position of a lineage in the phylogeny rather than on any explicit representation of interactions between residues.

The contribution of this study is a conditioning strategy rather than a general-purpose predictor. The improvement in prompt dependence over a sequence-only model, while consistent across both prediction tasks, is moderate in absolute terms, and evoPLM-Tree is not intended to compete with large pretrained protein language models on arbitrary proteins. Instead we show that phylogenetic quantities that are routinely computed during genomic surveillance, but are normally discarded before sequences reach a language model, carry information that a model uses when it is given access to them. The same conditioning scheme can be applied wherever mutation-annotated phylogenies exist, and becomes more useful as surveillance data accumulate for other pathogens.

Although evoPLM-Tree was not designed to recover exact descendant sequences, it accurately reproduced aggregate mutational patterns observed during SARS-CoV-2 evolution. Strong correlations between observed and predicted positional mutation frequencies indicate that the model learned which regions of spike are evolutionarily more or less permissive to change, consistent with experimental evidence for strongly heterogeneous mutational constraints across both the RBD and the broader spike protein [2–4]. Exact mutation recovery decreased with increasing evolutionary distance, possibly reflecting the growing number of plausible evolutionary trajectories separating ancestral and descendant sequences. This distinction highlights an important property of conditional generative models: multiple descendant sequences may be evolutionarily compatible with the same ancestral background, making exact sequence prediction inherently more difficult than recovering population-level mutational tendencies.

The enrichment of model probability among mutations tolerated in DMS experiments further suggests that the probabilities evoPLM-Tree assigns to descendant sequences capture information related to experimentally measurable protein function. Notably, evoPLM-Tree was trained exclusively on phylogenetically inferred ancestor–descendant sequence pairs without access to functional measurements. The stronger enrichment observed for RBD expression than for ACE2 binding may reflect the importance of maintaining folding and expression across evolutionary backgrounds, whereas ACE2 affinity represents only one component of spike fitness and can trade off with other selective pressures, including antibody escape [2, 3, 5, 25]. More broadly, experimental mutational measurements have been shown to contain information predictive of subsequent SARS-CoV-2 clade success [4], supporting the use of DMS as an external benchmark for evolutionary models.

Several limitations should be considered. The current study demonstrates evoPLM-Tree on SARS-CoV-2 spike evolution within the Omicron lineage. This choice reflects the data availability requirements of the approach: training and evaluating a model on ancestor–descendant sequence pairs requires a protein with dense genomic surveillance, a well-resolved phylogeny, and sufficient sampling across multiple lineages to construct a meaningful held-out set. SARS-CoV-2 spike is currently one of very few proteins for which all of these conditions are met, and it additionally benefits from rich existing knowledge of its evolutionary constraints and the availability of DMS data as an external benchmark. For most other viral proteins or organisms, equivalent data are not currently available, making cross-protein evaluation difficult. A further constraint is that only 53.5% of training-split records could be mapped to the 1 June 2022 mutation-annotated tree snapshot, compared with 99.4% for the held-out split. Training pairs may therefore be enriched for sequences deposited and placed on the tree early, and we cannot exclude that this introduces sampling bias relative to the full set of early Omicron genomes. The tree-derived features used here are also limited to phylogenetic topology and evolutionary distance, and the model does not incorporate lineage prevalence, geographic spread, or epidemiological growth rates. These distinctions are important because the probability a model assigns to a descendant sequence should not be interpreted as epidemiological fitness: a mutation may be compatible with a particular evolutionary trajectory without necessarily increasing lineage growth or prevalence. Furthermore, the training data represent inferred ancestor–descendant sequence pairs rather than directly observed transmission events, and descendant generation was evaluated primarily through aggregate mutational statistics rather than long-term forecasting of emerging variants. Whether similar performance can be achieved across other viral proteins, viruses, or broader evolutionary timescales remains an open question that future surveillance efforts may make tractable.

Future work could extend this framework in several directions. Incorporating representations from pretrained protein language models, which encode information related to protein structure, function, and mutational effects [14, 16, 17], may provide richer biochemical information while retaining explicit phylogenetic conditioning. More expressive tree representations or continuous phylogenetic embeddings could capture aspects of evolutionary history beyond the discretized features used here. Finally, evaluating evoPLM-Tree prospectively on newly emerging variants and integrating additional experimental measurements, including antibody escape and fitness assays, would provide a more comprehensive assessment of its utility for evolutionary forecasting. More broadly, as genomic surveillance expands to other pathogens and proteins, the ancestor–descendant sequence pairs needed to train and evaluate this type of model may become available beyond SARS-CoV-2. The exceptional sampling density of SARS-CoV-2 has made it possible to demonstrate that explicitly learning from phylogenetic sequence pairs is a viable and informative approach; this framework provides a template for applying the same idea wherever sufficiently dense and phylogenetically resolved sequence data exist.

## Materials and methods

### Phylogenetic data and sequence preprocessing

We downloaded human SARS-CoV-2 genomes (taxon 2697049) using the NCBI Datasets command-line tool [26]. We retained near-complete genomes of 28,500–31,000 nucleotides and processed them with Nextclade [11] to obtain quality-control annotations, Nextstrain clades, Pango lineages [27], and translated spike sequences. NCBI collection dates and geographic metadata were merged with the Nextclade results using accession identifiers.

We retained records with collection dates between 1 December 2019 and 31 December 2025; a Nextclade overall quality-control status other than “bad”; no reported processing errors or failed coding sequences; no “bad” private-mutation or SNP-cluster status; spike length between 1,260 and 1,285 amino acids; and at most three unknown residues (X) in spike. This procedure yielded 1,886,819 sequences. The receptor-binding domain (RBD) was defined as spike residues 319–541 inclusive (223 residues).

Training and held-out sets were separated temporally, with early Omicron lineages used for training and later Omicron lineages held out. Recombinant lineages, operationally defined as Pango designations beginning with X, were excluded. Training data comprised sequences from Nextstrain clades 21K, 21L, and 21M sampled before 1 June 2022 (265,674 records). The held-out dataset comprised non-recombinant sequences sampled on or after 1 June 2022 from clades 21K, 21L, 21M, 22A–22E, 23C, 23I, 24A–24C, 24E, 24G–24I, and 25A (451,933 records).

### Construction of ancestor–descendant sequence pairs from mutation-annotated trees

We used UShER mutation-annotated trees (MATs) [23] and their accompanying metadata to map NCBI accessions to MAT strain labels. To preserve the temporal separation, the training split was mapped to the 1 June 2022 public MAT snapshot, whereas the held-out split was mapped to the 21 January 2026 snapshot. The respective MAT match rates were 0.535 and 0.994. Only records with a resolved MAT leaf, MAT strain label, spike and RBD sequence, and valid collection date were eligible for pair construction.

For each MAT strain represented by multiple records, we retained one representative, prioritizing the largest duplicate count and then the latest collection date. We constructed a representative-leaf map by traversing upward from each retained leaf through at most 200 tree edges. At every traversed internal node, the closest observed descendant leaf was stored as that node’s representative.

For each candidate descendant leaf *v*, we considered its first, second, fourth, and eighth ancestors in the MAT. At each ancestor *a*, the mapped representative leaf *u* was used as the putative ancestral sequence, yielding an ordered sequence pair *u* → *v*. We retained at most two valid pairs per descendant, required *<* Δ_edges_ ≤200, and required non-decreasing root-to-tip nucleotide mutational pseudotime, *pt*_*u*_ ≤*pt*_*v*_. For each pair, we recorded the spike and RBD sequences, collection dates, Nextstrain clades and Pango lineages, tree depths, pseudotimes, hop count, path length in edges, and the root-to-tip nucleotide mutation difference Δ_nt_ = *pt*_*v*_ − *pt*_*u*_.

This yielded 282,158 training and 880,699 held-out raw sequence pairs. Before model training, we removed trivial sequence pairs (i.e., pairs with identical source and target sequences or no increase in root-to-tip nucleotide mutational pseudotime) separately for each prediction task: source and target spike/RBD sequences had to differ and Δ_nt_ ≥ 1. The resulting datasets contained 180,079/520,091 spike pairs and 87,370/272,133 RBD pairs for training/held-out evaluation, respectively.

### evoPLM-Tree architecture and training objective

evoPLM-Tree is a decoder-only causal language model implemented as a BERT language-model head with causal attention, in which each position can attend only to earlier positions [28, 29] (S1 Fig). The model has six transformer layers, hidden size 384, six attention heads, feed-forward size 1,536, and dropout probability The vocabulary contains 20 canonical amino acids, X, gap (-), special tokens, and discretized tree-feature tokens (11,810 tokens total). For each ancestor–descendant sequence pair *u* → *v*, the model input was

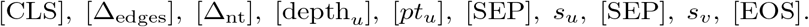

The four phylogenetic quantities were represented as bucket tokens. Tree-aware models therefore condition on the ancestral sequence and topology-derived features; sequence-only controls omitted the four tree tokens. During training, random spans of three tokens covering approximately 10% of the conditioning prefix were replaced by mask tokens. The target sequence was not masked.

The training labels were ignored for all prefix positions and retained only for the descendant sequence and [*EOS*]. Thus, the intended objective was to model the descendant sequence conditioned on the prefix:

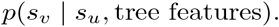

The loss was token-level causal cross-entropy computed only over the descendant sequence *s*_*v*_ and end-of-sequence token; all prefix tokens, including tree features and the source sequence, were excluded from the loss. Models were trained for three epochs with learning rate 3 *×* 10^−4^, weight decay 0.01, batch size 32, seed 0, and mixed-precision training when a GPU was available. Maximum context lengths were 1,024 tokens for RBD and 4,096 for full spike.

### Evaluation of conditioning dependence

To quantify reliance on conditioning information, models were evaluated under teacher forcing using the observed descendant sequences. For each sequence pair, the mean descendant-token cross-entropy was computed twice: once with the complete conditioning prompt available and once with the conditioning context masked from attention while leaving the input token identities unchanged. Masking was implemented by setting the attention-mask values of the conditioning tokens to zero.

For evaluation we used two model variants with the same transformer architecture: a sequence-only model and a tree-aware model (evoPLM-Tree). For the sequence-only model, the full input was

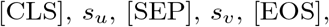

whereas for evoPLM-Tree it was

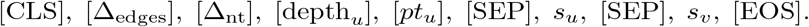

During the masked evaluation, all conditioning information (the ancestral sequence *s*_*u*_ and, for evoPLM-Tree, all tree-derived feature tokens) was removed from the attention context, while prediction targets remained unchanged. Conceptually, after masking the conditioning context, the model could attend to the following input. For the sequence-only model,

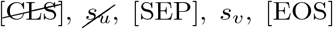

and for evoPLM-Tree,

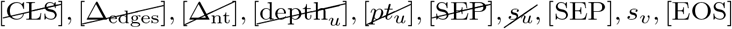

In both evaluations, the loss was computed only over the descendant sequence *s*_*v*_ (including the terminal [*EOS*] token), with all conditioning tokens excluded from the loss.

Prompt dependence was quantified as the ratio

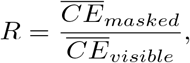

where 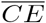 denotes the mean descendant-token cross-entropy. A ratio of 1 indicates that masking the conditioning prompt has no effect on prediction, whereas values greater than 1 indicate increasing reliance on conditioning information.

### Mutation frequency analysis

For analyses of positional mutation frequencies, descendant sequences were generated from 2,000 held-out sequence pairs sampled across quantile-defined strata of tree-edge distance (Δ_edges_) and mutational pseudotime difference (Δ_nt_), ensuring representation of both closely and distantly related sequence pairs. For each pair, we compared the input sequence with either the observed descendant sequence or a single model-generated descendant obtained by greedy decoding, in which the most probable amino acid was selected at each generation step. At each sequence position, mutation frequency was defined as the fraction of sequence pairs in which the descendant amino acid differed from the corresponding input amino acid, irrespective of the specific substitution.

Agreement between observed and predicted positional mutation-frequency profiles was quantified using Spearman’s rank correlation. Correlations and two-sided asymptotic p-values were computed with `scipy.stats.spearmanr (SciPy v1.15.2)` [30].

For structural visualization, positional mutation frequencies were mapped onto chain A of the SARS-CoV-2 spike structure from the Protein Data Bank [31] (PDB ID: 6VSB) by globally aligning the reference model sequence to the PDB sequence using a Needleman–Wunsch algorithm [32]. Only residues resolved in the experimental structure were assigned mutation-frequency values. The selected frequency profile was written to the B-factor field of the mapped residues and visualized in PyMOL v3.0.0 [33] using a common blue–white–red color scale for observed and predicted profiles. For receptor-binding domain (RBD) visualizations, only residues 319–541 were colored, whereas the remainder of the spike structure was displayed in gray.

### Deep mutational scanning enrichment analysis

To assess whether evoPLM-Tree preferentially assigns probability to experimentally tolerated mutations, we analyzed deep mutational scanning measurements for the Omicron BA.2 RBD obtained from MaveDB [34]. We used the score sets reported by Starr et al. [25], which quantify the effects of all single amino-acid substitutions on RBD expression and ACE2 binding (MaveDB IDs: `urn:mavedb:00000671-a-1` and `urn:mavedb:00000670-a-1`, respectively).

Each DMS dataset initially contained 4,020 amino-acid substitutions. Wild-type substitutions (n=201) and mutations with missing DMS measurements (n=18) were excluded, leaving 3,801 mutations for analysis in each dataset.

In the original study, ACE2-binding effects were derived from changes in − log_10_(*K*_*D*_), where *K*_*D*_ denotes the equilibrium dissociation constant between the RBD and ACE2, with lower values indicating stronger binding, and expression effects from changes in *log* mean fluorescence intensity, both relative to the parental BA.2 RBD. A score of zero therefore corresponds approximately to parental-like function, whereas positive values indicate increased ACE2 affinity or RBD expression. Mutations with DMS scores greater than or equal to zero were classified as tolerated, meaning that they performed at least as well as the parental BA.2 RBD for the measured phenotype.

For each mutation *m*, an indicator variable *T*_*m*_ was defined as

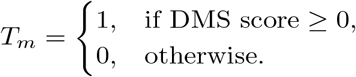

The probability-weighted fraction of tolerated mutations was calculated as

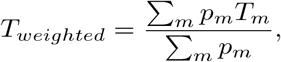

where *p*_*m*_ is the probability assigned by evoPLM-Tree to mutation *m*.

The background fraction of tolerated mutations was

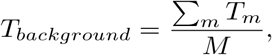

where *M* is the total number of analyzed mutations.

Weighted enrichment was then calculated as

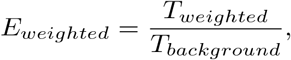

An enrichment of 1 corresponds to random allocation of probability mass across mutations, values greater than 1 indicate preferential assignment of probability to tolerated mutations, and values below 1 indicate preferential assignment to non-tolerated mutations.

### Figure generation

Figures used in this manuscript were generated using `Matplotlib v3.10.6` [35].

### Use of generative AI and AI-assisted technologies

During the preparation of this work, the authors used ChatGPT (OpenAI) and Claude (Anthropic) to refine sentence-level phrasing in the manuscript, particularly for non-native English speakers, and to assist with writing and debugging analysis code. All AI-suggested text was reviewed and edited by the authors. All AI-assisted code was inspected, tested, and verified by the authors against the reported results. No AI tool was used to generate, analyze, or modify the research data. All hypotheses, interpretations, results, conclusions, and limitations reported in this article are the authors’ own, and the authors take full responsibility for the content of the publication.

## Data availability

Code and reproducible workflows are available at GitHub, evoPLM-Tree (https://github.com/PlushZ/evoPLM-Tree). Source code (frozen release used in this study) is available at Zenodo, PlushZ/evoPLM-Tree: v1.0.0 (https://zenodo.org/records/22083791) [36]. Training and test datasets, model checkpoints are available at Zenodo (https://zenodo.org/records/22045909) [37].

## Author contributions statement

**Conceptualization:** PVP, WM, AFR. **Data curation:** PVP. **Formal analysis:** PVP. **Investigation:** PVP. **Methodology:** PVP, WM, AFR. **Project administration:** WM, AFR. **Software:** PVP. **Supervision:** WM, AFR. **Validation:** PVP. **Visualization:** PVP. **Writing – original draft:** PVP. **Writing – review & editing:** PVP, AFR.

## Acknowledgments

We thank Björn Grüning and the Freiburg Galaxy Team for computational infrastructure and support. We also thank Gavin Huttley for helpful discussions.

## Supporting information captions

**S1 Fig.**
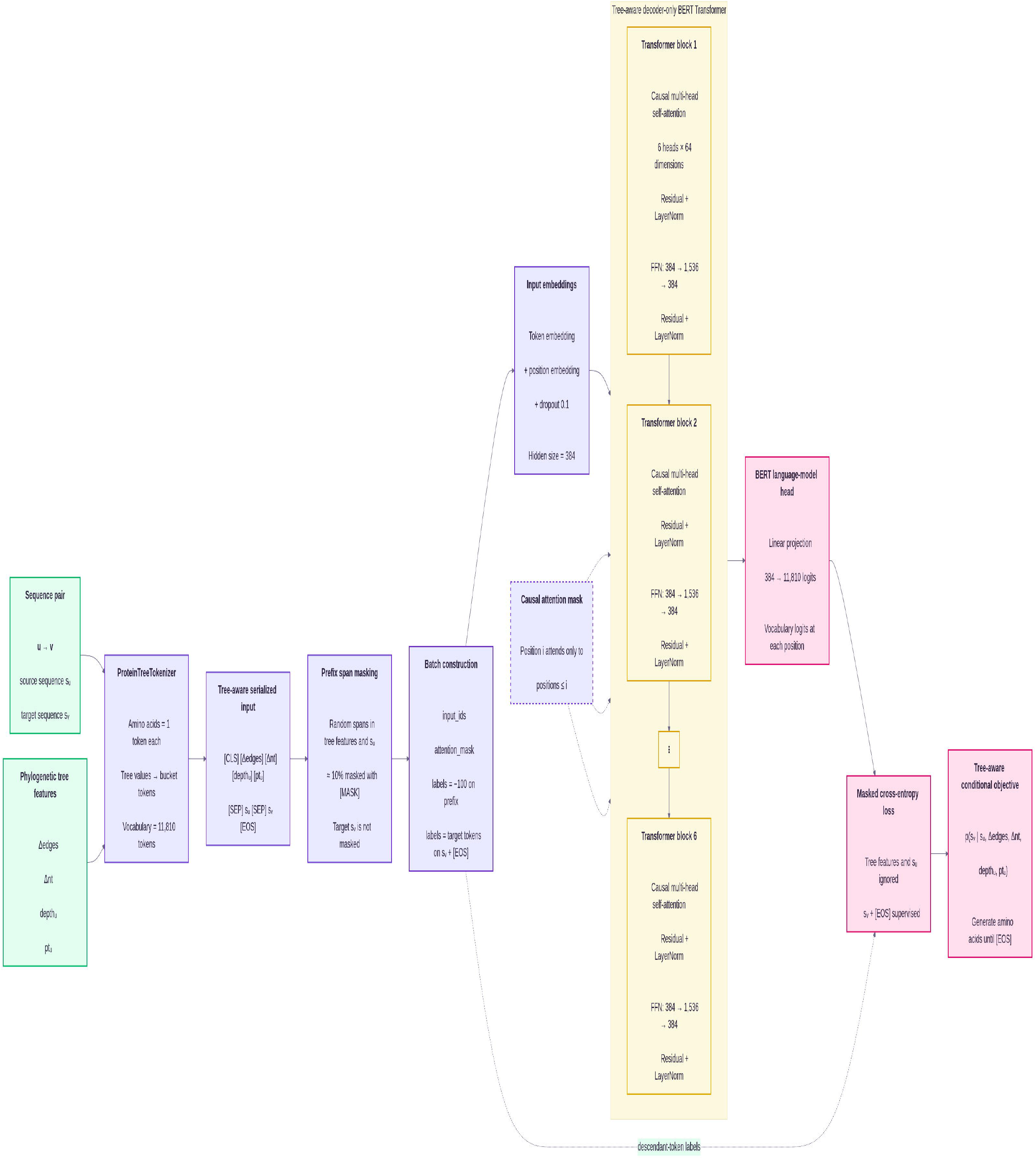
Architecture and training objective of the evoPLM-Tree model. Phylogenetic features and the ancestral protein sequence *s*_*u*_ are converted into a serialized token sequence and processed by a six-layer decoder-only Transformer with causal self-attention. During training, random spans covering approximately 10% of the conditioning prefix are replaced with [MASK] tokens, while the descendant sequence *s*_*v*_ remains unmasked and serves as the prediction target. Cross-entropy loss is computed only over the descendant sequence and terminal [EOS] token, with conditioning-prefix positions excluded from the loss. The model therefore learns the conditional distribution.

**S1 Table evoPLM-Tree generation results for the SARS-CoV-2 RBD**. Per-example results for 2,000 held-out sequence pairs, sampled with stratification by phylogenetic distance (`delta_edges`) and nucleotide mutation count (`delta_ nt_mutcount`; seed = 0). For each earlier sequence (`input_seq`), the model generated an RBD sequence (`pred_seq`), which was compared with the observed later sequence (true seq). Reported measures include sequence-level Hamming distance; observed, predicted, and correctly recovered mutation counts; and mutation precision, recall, and F1 score. A mutation was considered recovered only when its position and amino-acid substitution matched the observed mutation exactly. Additional columns provide phylogenetic, temporal, lineage, clade, and strain metadata for each sequence pair. Generation used greedy decoding conditioned on the input sequence and tree-feature tokens.

**S2 Table evoPLM-Tree generation results for the full-length SARS-CoV-2 spike protein**. Per-example results for 2,000 held-out sequence pairs, sampled with stratification by phylogenetic distance (`delta_edges`) and nucleotide mutation count (`delta_nt_mutcount`; seed = 0). For each earlier sequence (`input_seq`), the model generated a full-length spike sequence (`pred_seq`), which was compared with the observed later sequence (`true_seq`). Reported measures include sequence-level Hamming distance; observed, predicted, and correctly recovered mutation counts; and mutation precision, recall, and F1 score. A mutation was considered recovered only when its position and amino-acid substitution matched the observed mutation exactly. Additional columns provide phylogenetic, temporal, lineage, clade, and strain metadata for each sequence pair. Generation used greedy decoding conditioned on the input sequence and tree-feature tokens.

**S3 Table Mapping of observed and evoPLM-Tree-predicted RBD mutation hotspots onto the SARS-CoV-2 spike structure**. Residue-level mutation counts and frequencies were calculated across 2,000 held-out RBD sequence pairs by comparing each input sequence with the observed later sequence (`true`) or model-generated sequence (`pred`). Frequencies equal the corresponding counts divided by 2,000. RBD positions were aligned to residues 319–541 of chain A in the prefusion spike structure PDB 6VSB. The table reports zero- and one-based model coordinates, reference amino acid, observed and predicted hotspot statistics, mapping status, and corresponding PDB residue identifiers and amino acids. Of 223 RBD positions, 174 were mapped to resolved 6VSB residues; positions without structural coordinates are marked as unmapped.

**S4 Table Mapping of observed and evoPLM-Tree-predicted spike mutation hotspots onto the SARS-CoV-2 spike structure**. Residue-level mutation counts and frequencies were calculated across 2,000 held-out full-length spike sequence pairs by comparing each input sequence with the observed later sequence (`true`) or model-generated sequence (`pred`). Frequencies equal the corresponding counts divided by 2,000. Model coordinates were mapped to chain A of the prefusion spike structure PDB 6VSB by global sequence alignment. The table reports zero- and one-based model coordinates, reference amino acid, observed and predicted hotspot statistics, mapping status, and corresponding PDB residue identifiers and amino acids. Of 1,269 model positions, 959 were mapped to resolved 6VSB residues; positions absent from the experimental structure are marked as unmapped.

**S5 Table evoPLM-Tree probabilities and experimental effects for RBD–ACE2 binding**. Predicted mutation probabilities from the tree-aware model were matched to RBD–ACE2 binding deep mutational scanning scores from MaveDB dataset `urn:mavedb:00000670-a-1`. The table contains 3,801 non-wild-type substitutions with valid model probabilities and experimental scores. Columns report the assay-relative site, wild-type and substituted amino acids, predicted probability (prob), experimental DMS score, and binary tolerance classification (`dms_score` ≥ 0). Wild-type-equivalent substitutions were excluded; 18 additional substitutions at site 62 were omitted because their DMS scores were unavailable.

**S6 Table evoPLM-Tree probabilities and experimental effects for RBD expression**. Predicted mutation probabilities from the tree-aware model were matched to RBD expression deep mutational scanning scores from MaveDB dataset `urn:mavedb:00000671-a-1`. The table contains 3,801 non-wild-type substitutions with valid model probabilities and experimental scores. Columns report the assay-relative site, wild-type and substituted amino acids, predicted probability (`prob`), experimental DMS score, and binary tolerance classification (`dms_score` ≥ 0). Wild-type-equivalent substitutions were excluded; 18 additional substitutions at site 62 were omitted because their DMS scores were unavailable.

**S7 Table Summary of evoPLM-Tree performance across DMS datasets**. Performance was evaluated across two MaveDB datasets. For each dataset, the table reports the number of matched non-wild-type substitutions, Spearman correlation between predicted mutation probability and experimental DMS score with its nominal two-sided *P* value, and statistics for the three highest-probability substitutions per site. Top-*k* statistics include the number and mean DMS score of selected substitutions, the fractions classified as tolerated among selected and background substitutions, and their enrichment ratio. Probability-weighted tolerance is the predicted-probability-weighted mean of the binary tolerance indicator; weighted enrichment is its ratio to the unweighted background tolerance fraction. Substitutions were classified as tolerated when dms score ≥ 0.

